# The SEM-4 transcription factor is required for regulation of the oxidative stress response in *Caenorhabditis elegans*

**DOI:** 10.1101/2020.04.22.055772

**Authors:** Adilya Rafikova, Queenie Hu, Terrance J. Kubiseski

**Affiliations:** Department of Biology, York University, Toronto, Ontario, Canada M3J 1P3; Program in Neuroscience,’ York University, Toronto, Ontario, Canada M3J 1P3

**Author notes:** To whom correspondence should be addressed: Terrance Kubiseski, Department of Biology, York University, 4700 Keele Street, Toronto, ON, Canada, M3J 1P3.

**Keywords:** *Caenorhabditis elegans (C. elegans)*, aging, oxidative stress, genetics, gene expression, transcription factor, SEM-4, Splat-like (SALL) family of transcription factors

## Abstract

Oxidative stress causes damage to cells by creating reactive oxygen species (ROS) and the overproduction of ROS have been linked to the onset of premature ageing. We previously found that a *brap-2* (BRCA1 associated protein 2) mutant significantly increases the expression of phase II detoxification enzymes in *C. elegans.* An RNAi suppression screen to identify transcription factors involved in the production of *gst-4* mRNA in *brap-2* worms identified SEM-4 as a potential candidate. Here, we show that knockdown of *sem-4* suppresses the activation of *gst-4* caused by the mutation in *brap-2.* We also demonstrate that *sem-4 is* required for survival upon exposure to oxidative stress and that SEM-4 is required for expression of the transcription factor SKN-1C. These findings identify a novel role for SEM-4 in ROS detoxification by regulating expression of SKN-1C and the phase II detoxification genes.

**Article Summary:** Reactive oxygen species have been implicated as a harmful agent in many age-related diseases as well as an important signaling molecule. The transcription factor SKN-1 in C. elegans is an important regulator of reactive oxygen species levels. Here we show that the transcription factor SEM-4 is required to activate the expression of skn-1c and promote phase II detoxification gene expression. These findings identify a novel role for SEM-4 in regulating reactive oxygen species levels.

## Introduction

The nematode *C. elegans* has well-defined stress defense systems for protection from toxic compounds (1) and it offers a suitable model to dissect the gene regulatory network involved in the expression of stress response genes. Attention has been given to the transcription factors DAF-16/FOXO and SKN-1/Nrf2 due to their roles in regulating transcription in response to oxidative stress and lifespan extension in *C. elegans* (2–5). These factors regulate the transcription of detoxification genes such as *sod-3* and *gst-4,* to regulate levels of ROS (6–8). Although our understanding of the oxidative stress response has improved, the identification of transcription factors involved in this process and the control of their activities remain to be fully characterized.

Mammalian Brap2 (<u>Br</u>ca1 <u>a</u>ssociated binding <u>p</u>rotein 2; Brap as listed in the HUGO database) is a Ras-responsive E3 ubiquitin ligase that functions as a modulator of the Ras signaling pathway by facilitating activation of the Erk upon cell stimulation (9–11) and also appears to act as a cytoplasmic retention protein for a number of proteins (12–15). We have shown before that the *C. elegans* homolog *brap-2* loss of function mutant increases the MAP kinase activity of PMK-1 that leads to the increased expression of a wide range of phase II detoxification genes such as the glutathione S-transferases *gst-4, gst-7, gst-10,* and *gsto-2,* the dehydrogenase *dhs-8,* the gamma glutamylcysteine synthetase *gcs-1* and the UDP-glucurononsyl transferase *ugt-13* (16). We also performed an RNAi screen and showed that BRAP-2 regulates the transcription factor SKN-1 for the induction of phase II detoxification genes. SKN-1 is the *C. elegans* Nrf2 ortholog and, along with DAF-16, is a major transcriptional regulator of stress response in nematodes (5). Here we investigate the possibility that SEM-4, another candidate from the RNAi screen, is required for the oxidative stress response in the *brap-2* is mutant and during oxidative conditions.

SEM-4 (Sex Muscle abnormal 4) is a zinc finger (ZnF) containing transcription factor in *C. elegans* that is involved in neuronal, vulval and body wall muscle cell fate (17). The *sem-4* mutant worms are egg laying defective due to the transformation of sex myoblasts to body muscle cells. In addition to the development of a proper vulva, expression of SEM-4 has been shown to be important for the proper development and function of motor neurons and touch receptor neurons in the animal (17–19). SEM-4 is part of the NODE complex (CEH-6, EGL-27, SOX-2 (SRY (sex determining region Y)-box 2) and SEM-4) that activates EGL-5, and allows transformation of the Y cell (rectal epithelial cell) into PDA cell (motor neuron) (19).

Here, we report that SEM-4 has novel role in the SKN-1 dependent oxidative stress response as it promotes expression of phase II detoxification genes, likely through regulating *skn-1c* gene expression, and we demonstrate that *sem-4 is* required for survival upon exposure to oxidative stress. This work provides evidence that SEM-4 is part of the complex regulatory network that controls expression of genes involved in the oxidative stress response.

## Methods and Materials

### C. elegans Strains

All *C. elegans* strains were maintained as described by Brenner (20). Double mutant strains were generated according to standard protocols. Unless stated otherwise, worm strains were provided by the *Caenorhabditis* Genetics Center (CGC, University of Minnesota). Strains used in this study are listed in Table S1.

### Fluorescence microscopy

L4 *gst-4p::gfp* expressing worms were anesthetized using 2 mM Levamisole (Sigma L9756) and mounted on 2% agarose pad. Images of fluorescent worms were taken using a Zeiss LSM 700 confocal laser-scanning microscope with Zen 2010 Software.

### RNA isolation and Quantitative Real Time PCR

Worm RNA isolation, quantitative RT-PCR and data analysis were executed as described previously (16). Quantitative RT-PCR data were derived from 3 independent biological replicates and were analyzed using the comparative method (ΔΔCt). Results were graphed, and the relative expression of each strain was compared to N2. The endogenous control used for normalization was *act-1.* Primer sequences were as described previously (16).

### Oxidative stress assays and survival

Paraquat (Methyl viologen dichloride hydrate; Sigma (856177 Aldrich)) was dissolved in H2O, kept at −20°C and used as required. Strains were grown to the L4 stage before treatment. In experiments involving ROS and gene expression quantification, worms were placed in 100 mM paraquat for 1 hour, and then washed with M9 buffer three times. For survival assays, L4 stage worms were transferred to plates containing 2 mM paraquat and scored for survival every day. Worms were transferred to fresh plates every five days. All experiments were performed in three independent replicates.

### ROS quantification assay

The production of ROS (reactive oxygen species) was quantified using, 2,7-dichlorodihydrofluorescein-diacetate (DCFDA) (21). The levels of ROS were measured before and after paraquat exposure using a hybrid multimode microplate reader. All strains were synchronized and grown to L4 stage and then incubated with 0 mM or 100 mM paraquat for 1 hour. After incubation, worms were washed with M9 buffer three times, 200 worms/well from each strain were transferred into each well of a black 96-well plate and mixed with 100 μL of 50 μM DCFDA (diluted in 1X PBS). The fluorescence was measured kinetically every two minutes for 200 minutes using a BioTek, Synergy H4 microplate reader at excitation 485 nm and emission 520 nm, at 25°C.

### Statistics

Statistical analysis was performed using Prism 7 or 8 software (GraphPad). Statistical significance was determined using an unpaired student’s t-test when two means were compared and corrected for multiple comparisons using the Holm-Sidak method. P values of <0.05 were taken to indicate statistical significance. Error bars represent +/− standard error of the mean.

## Results

### The transcription factor SEM-4 is required for phase II detoxification gene expression in brap-2(ok1492) animals and upon oxidative stress

The goal of our lab is to further understand the molecular network of the phase II oxidative stress response. Previously, an RNAi screen identified 18 candidate transcription factors that were required for the enhanced expression of the phase II detoxification gene *gst-4* in the presence of the *brap-2(ok1492)* mutation (16). One of the candidates was SEM-4 and we set out to validate its role in activating *gst-4* expression. We obtained two strains that carry different mutant alleles of *sem-4, sem-4(n1378)* (which contains a C-terminal mutation) and *sem-4(n1971)* (which contains an N-terminal mutation). We generated *sem-4(n1378); brap-2(ok1492)* and *sem-4(n1971); brap-2(ok1492)* mutant worms containing the *gst-4p::gfp* transgene and found that the strains containing the *sem-4* mutations showed a weaker GFP expression compared to the *brap-2* single mutant (Fig 1A and 1B).

**Figure 1:**
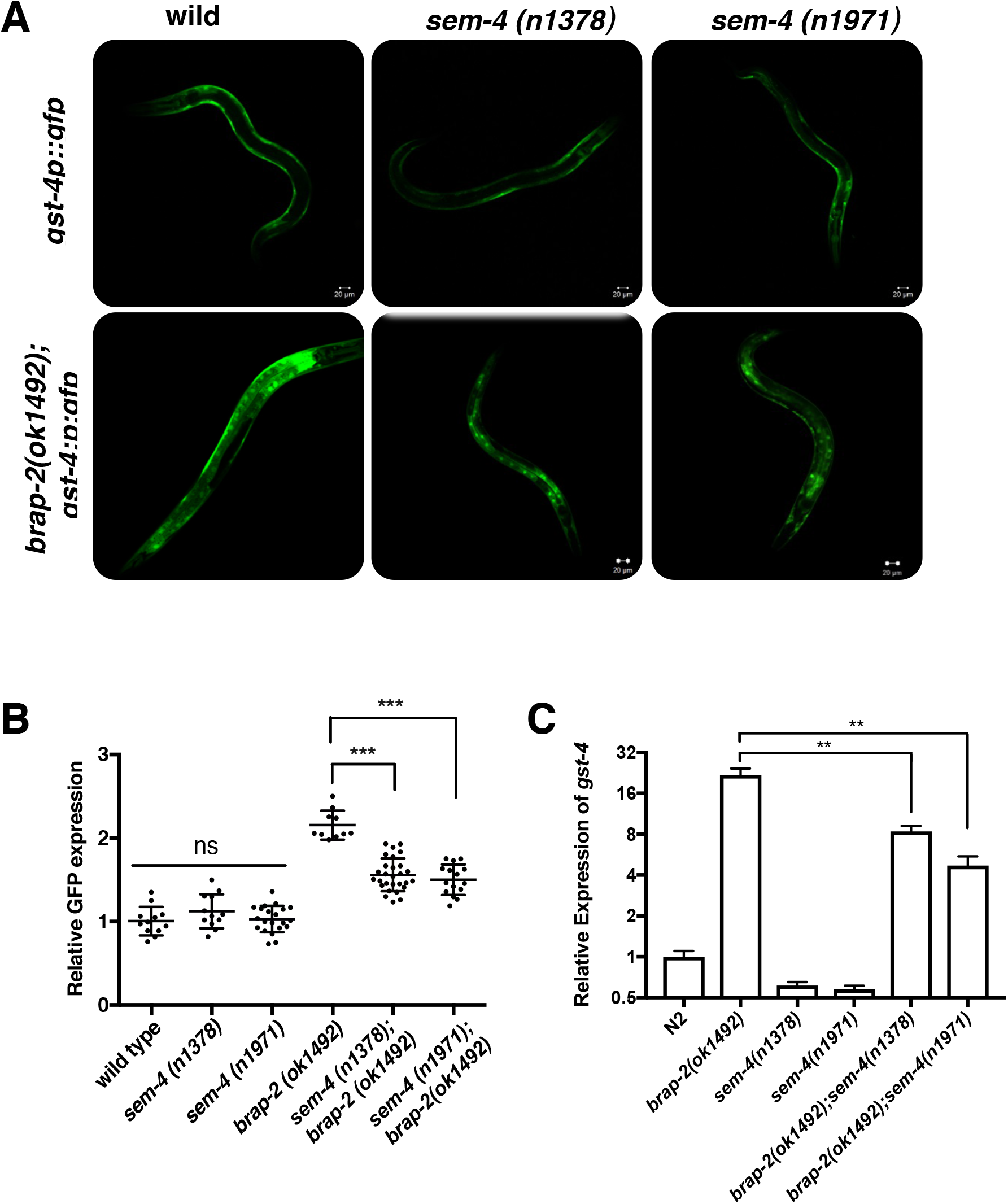
*gst-4* expression is reduced in *sem-4 mutant* worms. (A) GFP expression in *brap-2(ok1492); gst-4::gfp*, *sem-4(n1378); brap-2(ok1492); gst-4::gfp* and *sem-4(n1971); brap-2(ok1492); gst-4::gfp* mutant worms. (B). Images were analyzed with ImageJ software and resulted in significant difference in GFP expression. GFP intensity is lower in both *sem-4* mutant worms (n values between 12 and 28). (C) The *gst-4* mRNA was quantified by quantitative RT-PCR in *sem-4; brap-2* mutant worms show significant decrease in comparison to *brap-2* mutant worms. p<0.001***, p<0.01**.

We also measured *gst-4* mRNA by quantitative RT-PCR for the both of the *sem-4; brap-2* double mutant strains and found a significant reduction compared to the *brap-2* single mutant (Fig 1C). Other phase II detoxification genes *(gst-7, gst-10* and *gcs-1*) were assayed and we found that SEM-4 was required for the enhanced levels of expression in *brap-2(ok1492)* for *gst-7* and *gcs-1,* while the *sem-4* mutants did not significantly alter the expression of *gst-10* (Fig 2), possibly due to the existence of other regulating factors specific for *gst-10*.

**Figure 2:**
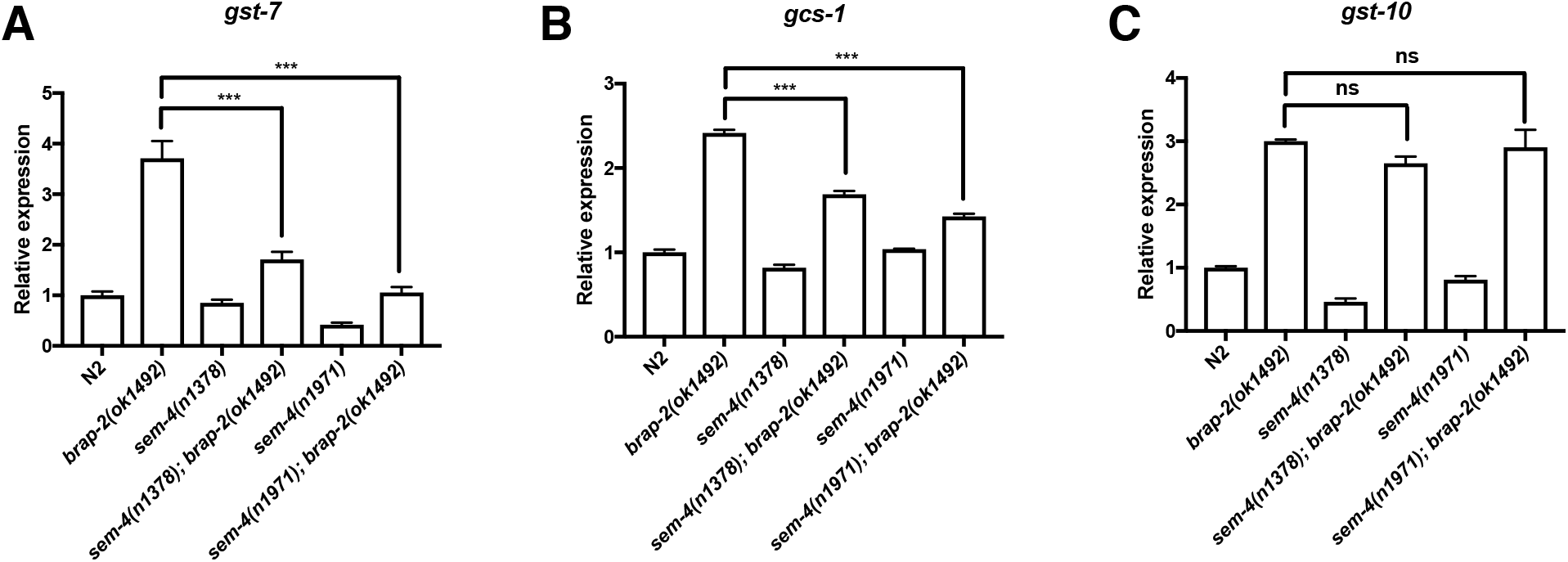
SEM-4 is required to promote phase II detoxification gene expression in *brap-2(ok1492)* worms. The mRNA levels of (A) *gst-7,* (B) *gcs-1* and (C) *gst-10* were quantified using quantitative RT-PCR. The *sem-4(n1378); brap-2(ok1492)* double mutant showed a reduction of mRNA expression for each gene tested. p<0.001***.

We also assessed the contribution SEM-4 plays during oxidative stress. To do this, we transferred the *gst-4p::gfp* transgene into *sem-4(n1378)* and *sem-4(n1971)* single mutants, exposed the worms to paraquat (which induces superoxide formation) and compared their fluorescent levels to wild type worms. We found that in the absence of paraquat, there is no significant difference in *gst-4* promoter activity between wild type and *sem-4* mutants. However, upon paraquat exposure, wild type worms demonstrated an increase in GFP levels that were significantly reduced in the presence of either the *sem-4(n1378)* or *sem-4(n1971)* alleles (Fig 3). Together, these results indicate that functional SEM-4 is required for the induction of phase II detoxification enzymes in the absence of BRAP-2 and upon exposure to oxidative stress.

**Figure 3:**
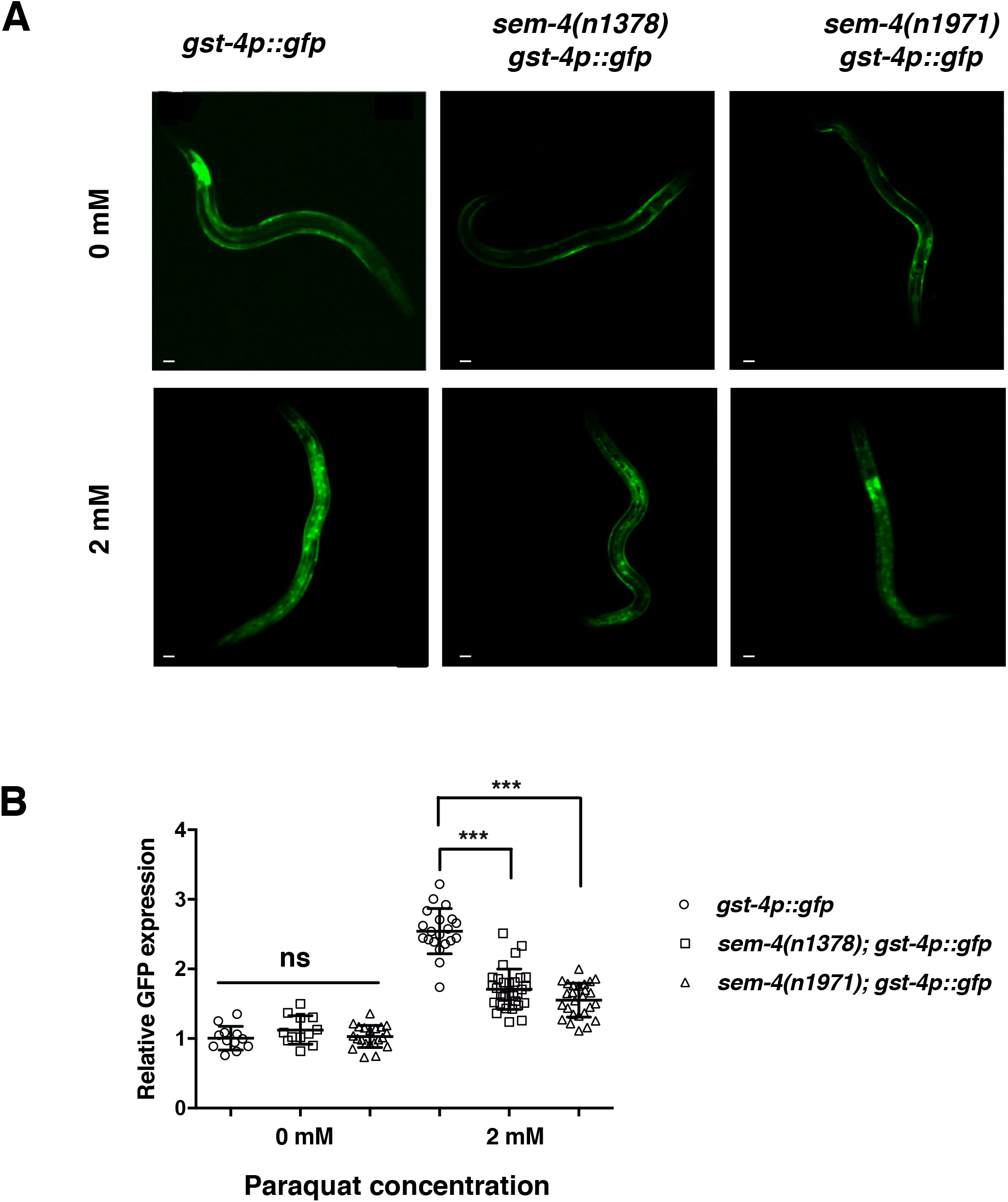
SEM-4 is required to promote *gst-4::gfp* expression during oxidative stress. (A) Fluorescent images of *gst-4p::gfp, sem-4(n1378); gst-4p::gfp* and *sem-4(n1971); gst-4::gfp* worms grown in the absence and presence of 2 mM paraquat. (B) Images were analyzed with ImageJ software and GFP intensity is lower in *sem-4* mutant worms grown on paraquat compared to wild type worms (n values between 12 and 31). p<0.001***, p<0.01**, p<0.05*.

### SEM-4 regulates expression of skn-1c

To assess if SEM-4 directly interacts with phase II detoxification genes, we surveyed the modENCODE genome-wide ChIP data (22). We found that for the five phase II detoxification genes that we surveyed *(gst-4, gcs-1, gsto-2, dhs-8* and *sdz-8)* that, under basal conditions, there were low levels of interaction of SEM-4 (ChIP-seq signal density scores between 10 and 30; Fig S1) to the promoter regions of these genes. Alternatively, we found a relatively higher level of interaction of SEM-4 to *skn-1* (two peaks with a density score of ~100), suggesting that SEM-4 may influence phase II detoxification gene expression indirectly through regulating *skn-1* gene expression rather than directly interacting with phase II detoxification gene promoters.

*C. elegans skn-1* encodes at least three splice variants (SKN-1A, SKN-1B and SKN-1C), each with its own distinct expression pattern and function. SKN-1A is an ER associated isoform involved in promoting transcriptional response to proteasomal dysfunction while SKN-1B and SKN-1C have roles in caloric restriction and stress resistance, respectively (4, 23, 24). Since SKN-1C is the isoform responsible for regulating phase II detoxification genes in *C. elegans,* we sought to determine if SEM-4 played a role in regulating *skn-1c* expression levels by quantifying *skn-1c* mRNA levels by quantitative RT-PCR. We found that under basal conditions, the expression of *skn-1c* decreased by 40-50% in *sem-4* mutant worms, while a transgenic strain for expression of SEM-4 (OP57 strain referred to as *sem-4(+))* showed a 30% increase in *skn-1c* mRNA levels (Fig 4A). Since we previously showed that *brap-2(ok1492)* mutants have higher levels of *skn-1* mRNA than wild type animals (16), we asked if *sem-4* is required for this increased *skn-1c* expression. We measured the mRNA levels of *skn-1c* in the two *sem-4; brap-2* strains and found a decrease in the amount of *skn-1* mRNA compared to the *brap-2* single mutant, restoring it to wild type levels (Fig 4B). This indicates that SEM-4 probably plays a role inducing *skn-1c* expression and that the decrease in expression of phase II detoxification genes in *sem-4; brap-2* double mutants may be due to the decrease in *skn-1* mRNA levels.

**Figure 4:**
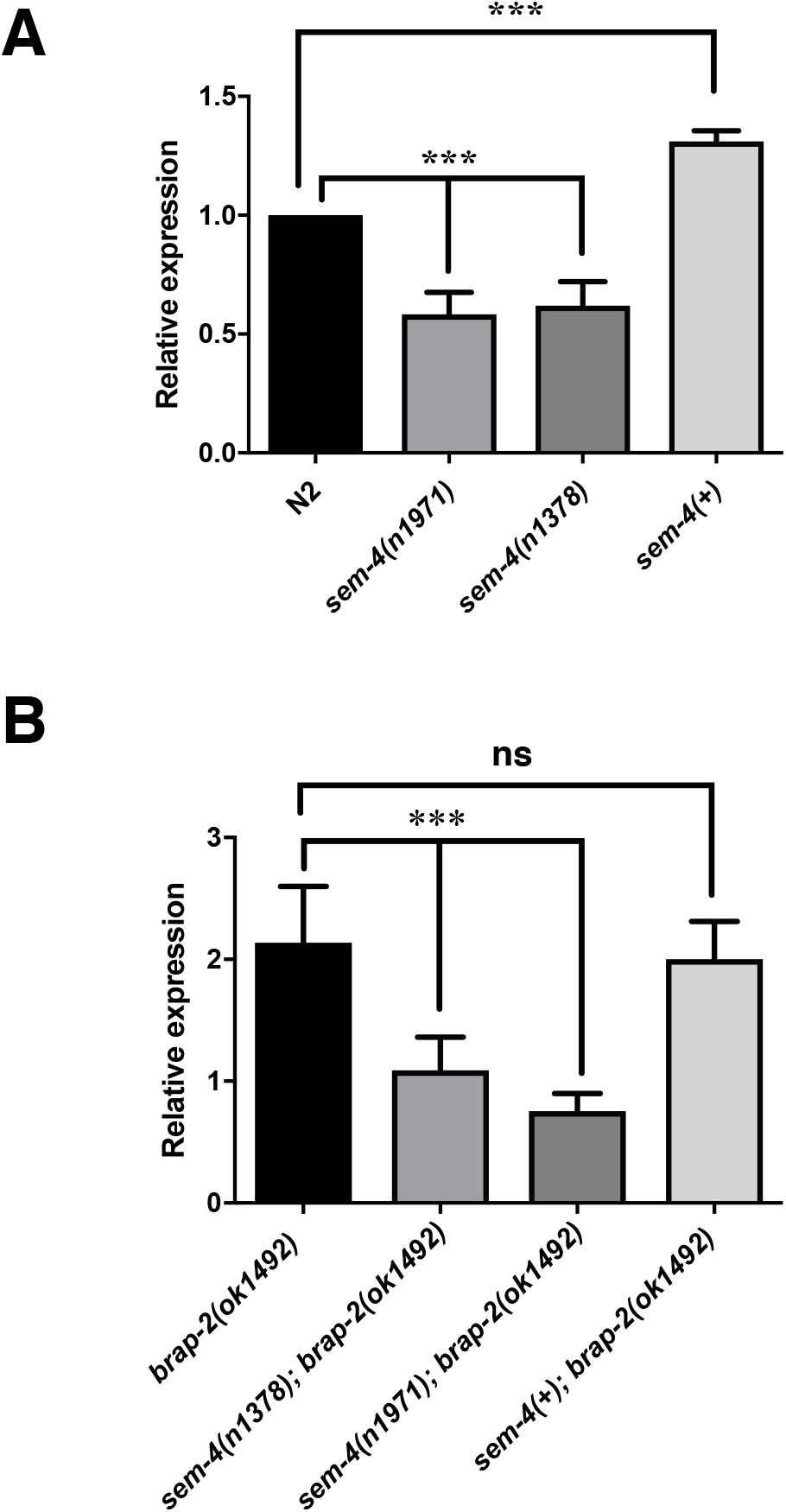
SEM-4 regulates the expression of *skn-1c.* (A) mRNA levels of *skn-1c* are affected by SEM-4. Overexpression of *sem-4* lead to increase of *skn-1c* mRNA by 30% (p<0.001, n=3 trials), while *sem-4(n1378)* mutant showed a decrease of 40% (p<0.001, n=5) and *sem-4(n1971)* by 50% (p<0.001, n=3). (B) SEM-4 regulated expression of *skn-1c* in *brap-2* strain. Significantly lower level of *skn-1c* mRNA is reported in *sem-4(n1971); brap-2* (65% decrease, p<0.001) and *sem-4(n1378); brap-2* (54% decrease, p<0.001) double mutants compared to *brap-2* worms. mRNA levels were quantified by quantitative RT-PCR with *act-1* as a reference gene. p<0.001***.

### sem-4 is required to reduce ROS levels in vivo and promote survival upon oxidative stress

We also used a functional assay to determine the ability of SEM-4 to increase phase II detoxification enzyme levels and reduce ROS *in vivo.* To do this, we quantified ROS levels using 2,7-dichlorodihydrofluoresceindiacetate (DCFDA) dye in three strains: N2, *sem-4(n1378),* and *sem-4(n1971)* upon exposure to 0 or 100 mM paraquat. As expected, we observed a significant increase ROS production in both *sem-4* strains in comparison to N2 (Fig 5). Therefore, the loss of *sem-4* elevates ROS production following oxidative stress induction, demonstrating its importance in regulating the detoxification of ROS.

**Figure 5:**
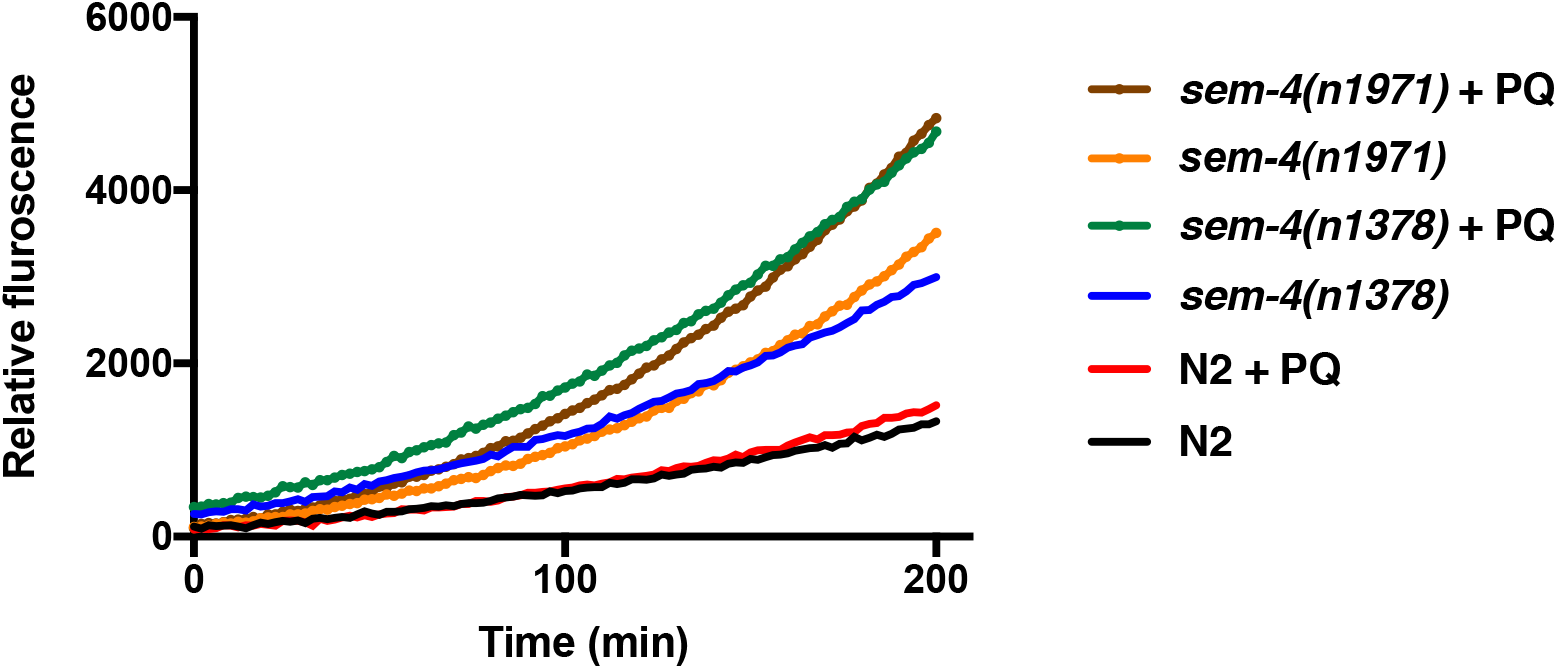
SEM-4 is required to reduce ROS production *in vivo*. Wild type and *sem-4* mutant strains were treated with 0 mM or 100 mM paraquat (+PQ) for 1 h and then mixed with 50 μM DCFDA dye in 96-well plates and the fluorescent levels were read for 200 min. ROS production was recorded as relative fluorescence units (RFU). Both *sem-4 (n1378)* and *sem-4(n1971)* showed an increase in ROS production after paraquat treatment over 200 min compared to wild type (N2).

To test the importance of SEM-4 on the survival of worms following oxidative stress, we cultured *sem-4* mutant nematodes on plates containing paraquat. For this experiment, *sem-4(n1971), sem-4(n1378)* and N2 worms were grown to L4 stage and transferred to 2 mM paraquat + 0.05 mg/mL FUDR plates and observed daily (Fig 6). It was found that *sem-4(n1971)* worms die with a median survival of 79 hours, while the median survival for *sem-4(n1378)* and N2 were 239 hours and 363 hours respectively (Fig 6B). These results demonstrate that the loss of functional SEM-4 is detrimental for worm survival when exposed to paraquat and that the inability of *sem-4* worms to survive in oxidative stress conditions corresponds to the detection of very high levels of ROS *in vivo*.

**Figure 6:**
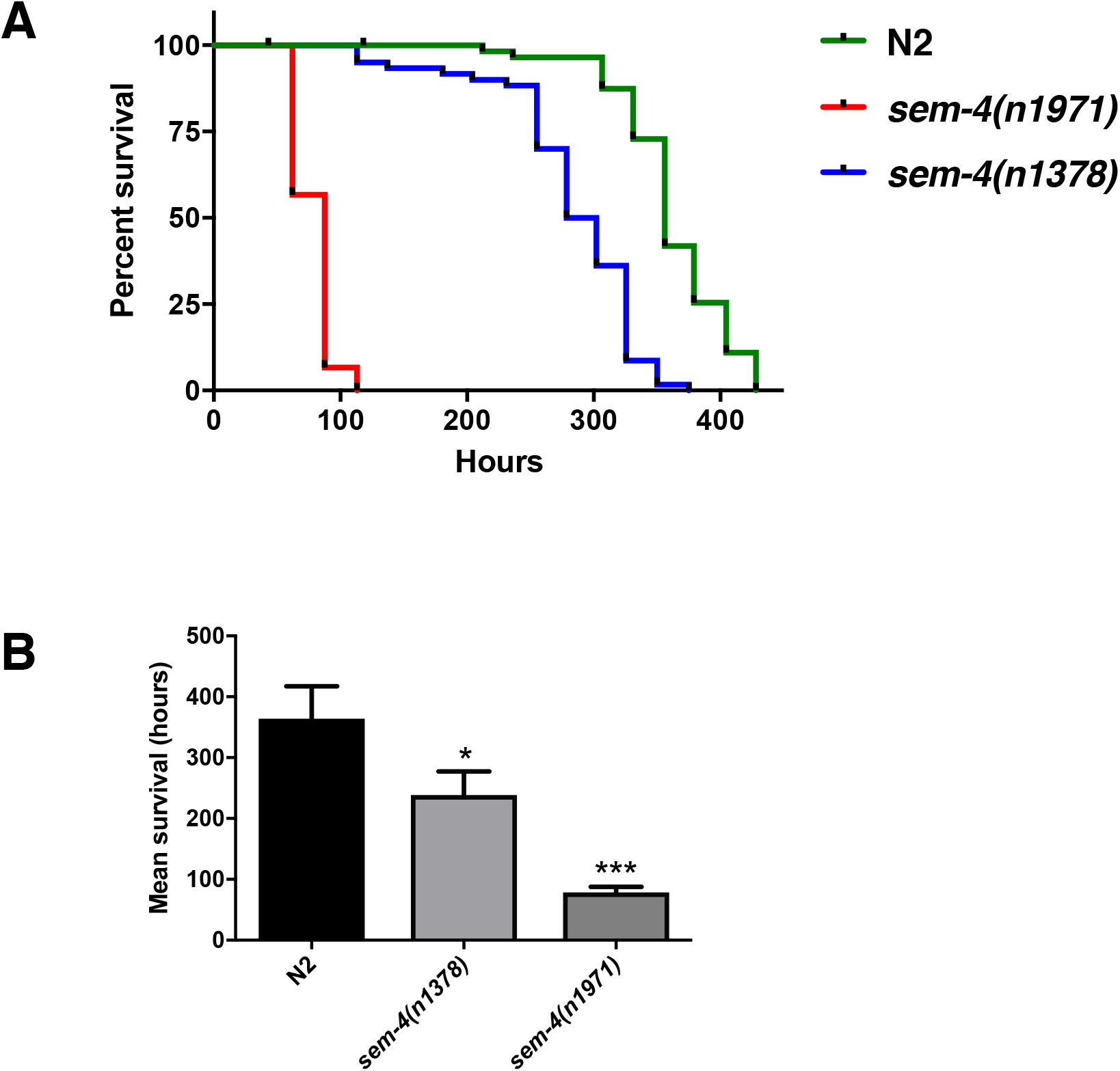
Survival following oxidative stress is dependent on SEM-4. (A) Survival of three strains were tested on 2 mM paraquat plates showed necessity of SEM-4 for *C. elegans.* The shortest lifespan is observed in *sem-4(n1971),* mean lifespan is 79 hours. The longest survival is observed in wild type worms, mean lifespan is 363 hours, and *sem-4(n1378)* worms survived for 283 hours. (B) Mean results for three trials. Survival of *sem-4(n1971)* is decreased by 78% (p<0.001) in comparison to N2 and survival of *sem-4(n1378)* is decreased by 34% (p=0.0303). Data was analyzed using OASIS software. p<0.001***, p<0.05*.

## Discussion

We have previously shown that BRAP-2 regulates the p38 MAP Kinase (PMK-1) in *C. elegans* (16). In the *brap-2* mutant and upon oxidative stress, activated PMK-1 phosphorylates SKN-1C to promote its re-localization from the cytosol to the nucleus where it promotes gene expression of phase II detoxification genes. Our work here indicates that the transcription factor SEM-4 is also required in response to oxidative stress to promote phase II gene expression, probably through the regulation of *skn-1c* gene expression, although we cannot rule out that SEM-4 may directly bind to phase II detoxification gene promoters. Like SKN-1, SEM-4 was initially discovered based on its essential role in development (17), but from these results it is apparent that SEM-4 also has functions that include maintaining cellular homeostasis in terminally differentiated cells. Although this study investigates SEM-4 role in phase II oxidative stress response, it remains to be determined if it is also involved in other stress response pathways.

This study also suggests that the mammalian orthologs of SEM-4 may be involved in this process. There are four known mammalian genes: SALL1/2/3/4 that share homology with SEM-4 and, to our knowledge, none of these gene products are associated with oxidative stress. There are known diseases associated with mutations in different SALL genes in humans. For instance, a mutation in SALL1 is linked to the autosomal disease, called Townes-Brocks syndrome (25), while mutant forms of SALL2 are present in human ovarian carcinoma (18) and a mutation in SALL4 leads to Okihiro syndrome (Duane-radial ray syndrome) (26). The function of SALL3 was recently discovered as an inhibitor of DNA methylation (27). Therefore, we anticipate that this study will provide a framework for further investigation of SALL1/2/3/4 in the mammalian oxidative stress response that may play a pathological role in a number of tissues and human diseases.

While the detrimental effects of ROS on the function of biological macromolecules is believed to lead to aging and cell death, ROS also acts as a fundamental signaling molecule in numerous processes, with hydrogen peroxide and singlet molecular oxygen having significant second messenger roles (28–31). The interplay between excessive oxidative conditions and maintenance of appropriate physiological levels of ROS requires the regulation of levels of detoxification enzymes by stress responsive transcription factors (32). Interestingly a number of transcription factors that have previously been demonstrated to play important developmental roles, such as *skn-1*, *elt-3* and *sem-4*, appear to have been repurposed in differentiated cells to function as regulators of ROS signalling. Further studies are required to understand how these (and other) factors coordinate together to properly regulate ROS levels.

## Acknowledgements

A number of strains were provided by the CGC, which is funded by NIH Office of Research Infrastructure Programs (P40 OD010440). A.R., Q.H. and T.J.K were supported by a grant from the Natural Sciences and Engineering Research Council of Canada.

## FOOTNOTE

The abbreviations used are: ROS, reactive oxygen species; FUDR, fluorodeoxyuridine; DCFDA, 2,7-dichlorodihydrofluorescein-diacetate; RFU, relative fluorescent units.

## References

1. Van Raamsdonk, J. M., and Hekimi, S. (2010) Reactive Oxygen Species and Aging in Caenorhabditis elegans: Causal or Casual Relationship? Antioxid. Redox Signal. 13, 1911–1953

2. Kenyon, C., Chang, J., Gensch, E., Rudner, A., and Tabtiang, R. (1993) A C. elegans mutant that lives twice as long as wild type. Nature. 366, 461–464

3. Murphy, C. T., McCarroll, S. A., Bargmann, C. I., Fraser, A., Kamath, R. S., Ahringer, J., Li, H., and Kenyon, C. (2003) Genes that act downstream of DAF-16 to influence the lifespan of Caenorhabditis elegans. Nature. 424, 277–283

4. An, J. H., and Blackwell, T. K. (2003) SKN-1 links C. elegans mesendodermal specification to a conserved oxidative stress response. Genes Dev. 17, 1882–1893

5. Blackwell, T. K., Steinbaugh, M. J., Hourihan, J. M., Ewald, C. Y., and Isik, M. (2015) SKN-1/Nrf, stress responses, and aging in Caenorhabditis elegans. Free Radic. Biol. Med. 88, 290–301

6. Oliveira, R. P., Porter Abate, J., Dilks, K., Landis, J., Ashraf, J., Murphy, C. T., and Blackwell, T. K. (2009) Condition-adapted stress and longevity gene regulation by Caenorhabditis elegans SKN-1/Nrf. Aging Cell. 8, 524–541

7. Wang, J., Robida-Stubbs, S., Tullet, J. M. A., Rual, J.-F., Vidal, M., and Blackwell, T. K. (2010) RNAi screening implicates a SKN-1-dependent transcriptional response in stress resistance and longevity deriving from translation inhibition. PLoS Genet. 10.1371/journal.pgen.1001048

8. Shore, D. E., and Ruvkun, G. (2013) A cytoprotective perspective on longevity regulation. Trends Cell Biol. 23, 409–420

9. Ory, S., and Morrison, D. K. (2004) Signal transduction: implications for Ras-dependent ERK signaling. Curr. Biol. CB. 14, R277–278

10. Matheny, S. A., and White, M. A. (2006) Ras-sensitive IMP modulation of the Raf/MEK/ERK cascade through KSR1. Methods Enzymol. 407, 237–247

11. Matheny, S. A., and White, M. A. (2009) Signaling threshold regulation by the Ras effector IMP. J. Biol. Chem. 284, 11007–11011

12. Li, S., Ku, C. Y., Farmer, A. A., Cong, Y. S., Chen, C. F., and Lee, W. H. (1998) Identification of a novel cytoplasmic protein that specifically binds to nuclear localization signal motifs. J. Biol. Chem. 273, 6183–6189

13. Asada, M., Ohmi, K., Delia, D., Enosawa, S., Suzuki, S., Yuo, A., Suzuki, H., and Mizutani, S. (2004) Brap2 functions as a cytoplasmic retention protein for p21 during monocyte differentiation. Mol. Cell. Biol. 24, 8236–8243

14. Chen, J., Hu, H., Zhang, S., He, M., and Hu, R. (2009) Brap2 facilitates HsCdc14A Lys-63 linked ubiquitin modification. Biotechnol Lett. 5, 615–21

15. Davies, R. G., Wagstaff, K. M., McLaughlin, E. A., Loveland, K. L., and Jans, D. A. (2013) The BRCA1-binding protein BRAP2 can act as a cytoplasmic retention factor for nuclear and nuclear envelope-localizing testicular proteins. Biochim. Biophys. Acta BBA – Mol. Cell Res. 1833, 3436–3444

16. Hu, Q., D’Amora, D. R., MacNeil, L. T., Walhout, A. J. M., and Kubiseski, T. J. (2017) The Oxidative Stress Response in Caenorhabditis elegans Requires the GATA Transcription Factor ELT-3 and SKN-1/Nrf2. Genetics. 206, 1909–1922

17. Basson, M., and Horvitz, H. R. (1996) The Caenorhabditis elegans gene sem-4 controls neuronal and mesodermal cell development and encodes a zinc finger protein. Genes Dev. 10, 1953–1965

18. Toker, A. S., Teng, Y., Ferreira, H. B., Emmons, S. W., and Chalfie, M. (2003) The Caenorhabditis elegans spalt-like gene sem-4 restricts touch cell fate by repressing the selector Hox gene egl-5 and the effector gene mec-3. Dev. Camb. Engl. 130, 3831–3840

19. Kagias, K., Ahier, A., Fischer, N., and Jarriault, S. (2012) Members of the NODE (Nanog and Oct4-associated deacetylase) complex and SOX-2 promote the initiation of a natural cellular reprogramming event in vivo. Proc. Natl. Acad. Sci. U. S. A. 109, 6596–6601

20. Brenner, S. (1974) The genetics of Caenorhabditis elegans. Genetics. 77, 71–94

21. Yang, H.-C., Chen, T.-L., Wu, Y.-H., Cheng, K.-P., Lin, Y.-H., Cheng, M.-L., Ho, H.-Y., Lo, S. J., and Chiu, D. T.-Y. (2013) Glucose 6-phosphate dehydrogenase deficiency enhances germ cell apoptosis and causes defective embryogenesis in Caenorhabditis elegans. Cell Death Dis. 4, e616

22. Gerstein, M. B., Lu, Z. J., Van Nostrand, E. L., Cheng, C., Arshinoff, B. I., Liu, T., Yip, K. Y., Robilotto, R., Rechtsteiner, A., Ikegami, K., Alves, P., Chateigner, A., Perry, M., Morris, M., Auerbach, R. K., Feng, X., Leng, J., Vielle, A., Niu, W., Rhrissorrakrai, K., Agarwal, A., Alexander, R. P., Barber, G., Brdlik, C. M., Brennan, J., Brouillet, J. J., Carr, A., Cheung, M.-S., Clawson, H., Contrino, S., Dannenberg, L. O., Dernburg, A. F., Desai, A., Dick, L., Dosé, A. C., Du, J., Egelhofer, T., Ercan, S., Euskirchen, G., Ewing, B., Feingold, E. A., Gassmann, R., Good, P. J., Green, P., Gullier, F., Gutwein, M., Guyer, M. S., Habegger, L., Han, T., Henikoff, J. G., Henz, S. R., Hinrichs, A., Holster, H., Hyman, T., Iniguez, A. L., Janette, J., Jensen, M., Kato, M., Kent, W. J., Kephart, E., Khivansara, V., Khurana, E., Kim, J. K., Kolasinska-Zwierz, P., Lai, E. C., Latorre, I., Leahey, A., Lewis, S., Lloyd, P., Lochovsky, L., Lowdon, R. F., Lubling, Y., Lyne, R., MacCoss, M., Mackowiak, S. D., Mangone, M., McKay, S., Mecenas, D., Merrihew, G., Miller, D. M., Muroyama, A., Murray, J. I., Ooi, S.-L., Pham, H., Phippen, T., Preston, E. A., Rajewsky, N., Rätsch, G., Rosenbaum, H., Rozowsky, J., Rutherford, K., Ruzanov, P., Sarov, M., Sasidharan, R., Sboner, A., Scheid, P., Segal, E., Shin, H., Shou, C., Slack, F. J., Slightam, C., Smith, R., Spencer, W. C., Stinson, E. O., Taing, S., Takasaki, T., Vafeados, D., Voronina, K., Wang, G., Washington, N. L., Whittle, C. M., Wu, B., Yan, K.-K., Zeller, G., Zha, Z., Zhong, M., Zhou, X., modENCODE Consortium, Ahringer, J., Strome, S., Gunsalus, K. C., Micklem, G., Liu, X. S., Reinke, V., Kim, S. K., Hillier, L. W., Henikoff, S., Piano, F., Snyder, M., Stein, L., Lieb, J. D., and Waterston, R. H. (2010) Integrative analysis of the Caenorhabditis elegans genome by the modENCODE project. Science. 330, 1775–1787

23. Bishop, N. A., and Guarente, L. (2007) Two neurons mediate diet-restriction-induced longevity in C. elegans. Nature. 447, 545–549

24. Glover-Cutter, K. M., Lin, S., and Blackwell, T. K. (2013) Integration of the unfolded protein and oxidative stress responses through SKN-1/Nrf PLoS Genet. 9, e1003701

25. Netzer, C., Bohlander, S. K., Hinzke, M., Chen, Y., and Kohlhase, J. (2006) Defining the heterochromatin localization and repression domains of SALL1. Biochim. Biophys. Acta. 1762, 386–391

26. Kohlhase, J., Chitayat, D., Kotzot, D., Ceylaner, S., Froster, U. G., Fuchs, S., Montgomery, T., and Rösler, B. (2005) SALL4 mutations in Okihiro syndrome (Duane-radial ray syndrome), acro-renal-ocular syndrome, and related disorders. Hum. Mutat. 26, 176–183

27. Shikauchi, Y., Saiura, A., Kubo, T., Niwa, Y., Yamamoto, J., Murase, Y., and Yoshikawa, H. (2009) SALL3 interacts with DNMT3A and shows the ability to inhibit CpG island methylation in hepatocellular carcinoma. Mol. Cell. Biol. 29, 1944–1958

28. Holmström, K. M., and Finkel, T. (2014) Cellular mechanisms and physiological consequences of redox-dependent signalling. Nat. Rev. Mol. Cell Biol. 15, 411–421

29. Shibata, Y., Branicky, R., Landaverde, I. O., and Hekimi, S. (2003) Redox regulation of germline and vulval development in Caenorhabditis elegans. Science. 302, 1779–1782

30. Putker, M., Madl, T., Vos, H. R., de Ruiter, H., Visscher, M., van den Berg, M. C. W., Kaplan, M., Korswagen, H. C., Boelens, R., Vermeulen, M., Burgering, B. M. T., and Dansen, T. B. (2013) Redox-dependent control of FOXO/DAF-16 by transportin-1. Mol. Cell. 49, 730–742

31. Sies, H., Berndt, C., and Jones, D. P. (2017) Oxidative Stress. Annu. Rev. Biochem. 86, 715–748

32. Sies, H., and Jones, D. P. (2020) Reactive oxygen species (ROS) as pleiotropic physiological signalling agents. Nat. Rev. Mol. Cell Biol. 10.1038/s41580-020-0230-3

